# A novel polymorphism in nitric oxide synthase interacting protein (NOSIP) modulates nitric oxide and mortality in Human Sepsis

**DOI:** 10.1101/038398

**Authors:** Ratnadeep Mukherjee, Diwakar Kumar Singh, Rajkumar Patra, Pijus Kanti Barman, Birendra Kumar Prusty, Pravat Thatoi, Rina Tripathy, Bidyut Kumar Das, Balachandran Ravindran

**Affiliations:** Infectious Disease Biology Group, Institute of Life Sciences, Bhubaneswar, INDIA; Department of Medicine, S.C.B. Medical College, Cuttack, INDIA; Post Graduate Department of Paediatrics, Sishu Bhawan, Cuttack, INDIA

**Author notes:** (BR), (RM). Equal contribution (Ratnadeep Mukherjee, Diwakar Kumar Singh).

## Abstract

Nitric oxide, synthesised by three isoforms of Nitric Oxide synthases viz., nNOS by neurons, eNOS by endothelial cells and iNOS by phagocytes, performs a wide variety of biological functions in neurons, vascular endothelial cells and immune cells. Interaction between inducible nitric oxide synthase (iNOS) and Nitric oxide synthase interacting protein (NOSIP) was observed both in human monocytes and mouse macrophages and in cell free systems by biophysical methods. A novel mutation in nitric oxide synthase interacting protein (NOSIP) determined NO levels produced by human monocytes and was associated with disease severity in Sepsis patients. The study reveals NOSIP as an important regulator of inflammation by virtue of its ability to influence nitric oxide production both in mice and in humans and opens up novel avenues for therapeutic strategies against acute inflammation. While the influence of this novel NOSIP polymorphism in cardio-vascular and neuronal functions could be a subject of future investigations, its role in determining disease severity and mortality of the ongoing Covid 19 pandemic will be of immediate relevance.

## Introduction

Nitric oxide (NO) is a well-known signalling molecule that modulates a wide variety of functions in endothelial cells [1], neurons [2] and immune cells [3, 4]. In the immune system its initial function was shown as a molecule with anti-microbial [5, 6] and tumoricidal [7, 8] functions. Later reports suggested NO to be a regulator of other facets of immune responses like inhibition of T and B lymphocyte proliferation [9–13] and leukocyte trafficking [14]; modulation of cytokine, chemokine and growth factor production [15–19]; T helper cell subtype differentiation [20–22] etc. The role of NO in regulating hyperinflammation has been a subject of contemporary interest since it appears to play a dual function of being inflammatory as well as antiinflammatory in different contexts.

Nitric oxide synthase (NOS), the enzyme that produces NO in mammalian cells using L-Arginine as its substrate [23], has a multitude of interacting partners that regulate its function, localization and trafficking [4, 24, 25]. Of the three NOS isoforms, NOS1 (nNOS) and NOS3 (eNOS) binding proteins have been fairly well characterized [24]. One such protein is nitric oxide synthase interacting protein (NOSIP), that has been reported to interact with both nNOS [26] and eNOS [27, 28] post-translationally and regulate their function. Surprisingly however the role of NOSIP and its interaction with NOS2 (iNOS) and its ability to regulate function of iNOS in immune cells has not been investigated so far and this study attempts to fill this lacuna through a combination of biochemical, biophysical and imaging techniques. More critically we provide evidence for the first time a novel mutation in a non-coding region of human NOSIP gene and its influence on NOSIP synthesis and association with enhanced mortality in human sepsis.

Many reports in literature describe links between NO and sepsis. Demonstration of elevated levels of NO by-products in sepsis [29] and observations on NOS inhibitors alleviating the hemodynamic manifestations of septic shock [30] led to the general belief that NO was detrimental to host during sepsis. This resulted in undertaking a randomized controlled trial that involved treatment of sepsis patients with a nonspecific NOS inhibitor [31]. However, the trial had to be truncated due to observed higher mortality in the drug treated group over placebo trated control group. On the other hand few reports do suggest a protective role for NO during inflammation and sepsis [32–34]. Using wild type and iNOS deficient mice models of acute inflammation and human sepsis, we demonstrate a protective role for nitric oxide during inflammation and sepsis. Finally, we show that species specific differences in nitric oxide levels produced by mouse and human monocytes can be attributed to differential expression of NOSIP.

## Materials and Methods

### Reagents and kits

Gram-negative bacterial lipopolysaccharide (LPS, E.coli O55:B5), Ficoll Histopaque, Blood Genomic DNA extraction kit, Duolink Proximity Ligation Assay (PLA) kit, Nitrite/Nitrate estimation kit, N-*ω*-Nitro-L-arginine methyl ester hydrochloride (L-NAME), Bovine Serum Albumin (BSA), Phorbol 12-myristate 13-acetate (PMA), Paraformaldehyde (PFA), iNOS recombinant protein, protease inhibitor, Isopropyl *β*-D-1-thiogalactopyranoside, Isopropyl *β*-D-thiogalactoside (IPTG), Ampicillin sod-ium salt, and Triton X-100 was obtained from Sigma chemicals. Dulbecco’s Modified Eagle’s Medium (DMEM), Foetal Bovine Serum (FBS), and Phosphate Buffered Saline (PBS) were from PAN Biotech GmbH. Acetic acid, citric acid, and dextrose were all from Fisher Scientific. Human 27-plex cytokine and Mouse 23-plex cytokine analysis kits, GLC sensor chip, amine coupling kit were procured from Bio-Rad Laboratories. FACS lysing solution, Lyse/Fix buffer, Permeabilization buffer IV and FACSComp beads were bought from BD Biosciences. Brefeldin A was obtained from eBiosciences. Taq DNA polymerase for PCR was from Thermo Scientific. 4’,6-Diamidino-2-Phenylindole, Dihydrochloride (DAPI) and ProLong® Gold Antifade were purchased from Molecular Probes. Nitric Oxide Synthase activity assay kit was purchased from Abcam. Ni-NTA agarose was purchased from Qiagen.

### Antibodies

Anti-goat IgG Alexa Fluor 488 and anti-rabbit IgG Alexa Fluor 647 antibodies were purchased from Molecular Probes. Rabbit polyclonal antibody to iNOS was obtained from Abcam and goat polyclonal antibody to NOSIP was purchased from SantaCruz Biotechnolgies.

### Human sepsis patients

The study was approved by ethics committees of Institute of Life Sciences and S.C.B. Medical College, and signed informed consent was obtained from all participants. 139 sepsis patients who were admitted to medical intensive care units at the Department of Medicine, S.C.B. Medical College and Hospital (Cuttack, India) were recruited for the study. For sepsis, patients were eligible for inclusion only if they had systemic inflammatory response syndrome and had a source of infection, proven or suspected. The acute physiological and chronic health evaluation II (APACHE II) scoring system was used to categorize the patients. Definitions of sepsis, severe sepsis, septic shock, and multi-organ dysfunction syndrome (MODS) were in accordance with published criteria [35, 36]. The following categories of patients were excluded from the study: patients with diabetes mellitus, hypertension, nephrotic syndrome, chronic kidney disease (sonographic feature of CKD and/or GFR*<*30 ml/min), patients with cardiac failure and immunocompromised individuals. Blood was collected in vials containing 15% (v/v) Acetate Citrate Dextrose (ACD). This was followed by isolation of plasma by centrifugation at 2000 rpm for 10 min. Plasma was stored in single-use aliquots at −80°C.

### Mouse model of endotoxemia

8-10 weeks old male C57BL/6 mice were used for the study. All animal experiments were approved by the institutional animal ethics committee of Institute of Life sciences, Bhubaneswar. To simulate two groups based on lethality in human sepsis cases, a 5mg/kg nonlethal dose and 35mg/kg lethal dose of gram negative bacterial lipopolysaccharide (E.coli O55:B5, Sigma) was injected intraperitoneally. The animals were sacrificed at 2, 4, 8 and 16 hours post injection to mimic early and late stages of endotoxemia. Blood was collected in vials containing ACD as anticoagulant (15% v/v). collected blood was centrifuged at 2000 rpm for 10 minutes for isolation of plasma. Isolated plasma was stored in 100 *µ*l single-use aliquots at −80°C.

### Assessment of nitric oxide deficiency on LPS – mediated inflammation

8 – 10 weeks old male BALB/C mice were injected with 100 mg/kg L-NAME once every 24 hours for inhibition of nitric oxide synthesis. To examine effect of nitric oxide synthase inhibition on LPS – mediated pathology, untreated or L-NAME treated mice were injected with 2 mg/kg LPS and mortality was monitored for 4 days. For comparing LPS toxicity between wild – type and *inos* knockout mice, male animals between 8-10 weeks age were injected with LPS at 15 mg/kg and either mortality was scored for 5 days or animals were sacrificed at 2, 6 and 12 hours to estimate cytokines in plasma.

### Ex vivo stimulation of human and mouse peripheral blood

Whole blood was withdrawn from apparently healthy human donors by venepuncture or from normal healthy mouse by cardiac puncture in acid citrate dextrose (ACD) anticoagulant at 15% v/v. Human donors were recruited from institute students. To test for cytokine expression, 50 *µ*l whole blood was left untreated or stimulated with LPS at 1 *µ*g/ml for 2 hours in 37°C water bath along with brefeldin A (eBiosciences) at 1:1000 dilution. Post stimulation, cells were stained with fluorochrome conjugated antibodies and analysed on a flow cytometer.

### Measurement of cytokines in plasma

Human plasma was analysed using the human 27-plex cytokine panel (Bio-Rad) according to manufacturer’s instructions and contained the following targets: IL-1*β*, IL-1ra, IL-2, IL-4, IL-5, IL-6, IL-7, IL-8, IL-9, IL-10, IL-12(p70), IL-13, IL-15, IL-17, Basic FGF, Eotaxin, G-CSF, GM-CSF, IFN-*γ*, IP-10, MCP-1, MIP-1*α*, MIP-1*β*, PDGF, RANTES, TNF-*α*, and VEGF. Mouse plasma was analysed using the mouse 23-plex cytokine panel (Bio-Rad) as specified by the manufacturer and contained the following targets: IL-1*α*, IL-1*β*, IL-2, IL-3, IL-4, IL-5, IL-6, IL-9, IL-10, IL-12(p40), IL-12(p70), IL-13, IL-17, Eotaxin, G-CSF, GM-CSF, IFN-*γ*, KC, MCP-1, MIP-1*α*, MIP-1*β*, RANTES, and TNF-*α*. All samples were read on a Bioplex 200 system (Bio-Rad). Concentrations of unknown samples were interpolated from a 8-point standard curve fitted with a five-parameter logistic regression model.

### Measurement of plasma nitrate and nitrite

Levels of nitrate and nitrite in plasma obtained from human and mice were estimated by a Griess colorimetric assay kit (Sigma chemicals) according to manufacturer’s instructions. Briefly, nitrate in plasma was first converted into nitrite by nitrate reductase. Total nitrite was then estimated by a colorimetric reaction involving Griess reagent A (a solution of sulphanilamide in phosphoric acid) and Griess reagent B (Napthylethylenediamine in phosphoric acid). Absorbance was read at 540 nm. Unknown concentrations were estimated from a standard curve generated with known concentrations of sodium nitrite.

### Assessment of intracellular localization of iNOS and NOSIP by confocal imaging

Mouse and human whole blood were fixed, lysed and permeabilized followed by staining with anti-iNOS and anti-NOSIP antibodies. Followed by washing, the cells were counterstained with DAPI, fixed on a slide and colocalization was scored on a Leica Laser Scanning Confocal Microscope. For studying interaction between iNOS and NOSIP, in situ Proximity Ligation Assay was used using a commercially available kit (Sigma Duolink) according to manufacturer’s instructions. In case of the human monocytic cell line THP-1, cells were grown in a coverslip - bottomed culture dish in complete IMDM media containing 10% fetal bovine serum. Cells were made to adhere by treatment with 10 nM Phorbol myristate acetate (PMA) for 48 hours. Following adherence, cells were washed with PBS and fixed with 2% paraformaldehyde (PFA). After rinsing with PBS, cells were permeabilized with 0.1% Triton X-100 following which they were incubated in a blocking buffer containing 2% bovine serum albumin (BSA) and 2% fetal bovine serum (FBS) in PBS to minimize nonspecific binding of antibodies. After blocking, cells were first incubated with primary antibody followed by fluorochrome conjugated secondary antibodies. Finally, cells were washed, counterstained with DAPI and image was acquired in a Leica SP5 confocal microscope.

### Imaging cytometry analysis of NOSIP expression

For comparing intracellular NOSIP expression between ins and del individuals, 100 *µ*l whole blood from each of 4 *ins* and 3 *del* individuals were fixed, lysed and permeabilized followed by staining with anti-NOSIP antibody. After washing, the cells were resuspended in PBS and acquired on an ImagestreamX imaging cytometer (Amnis corp.). 20,000 events per sample were acquired for analysis. The obtained events were gated and analysed by IDEAS 6.0 software.

### Cloning, Expression and Purification of iNOS and NOSIP proteins

The construct of human iNOS oxygenase domain (1284 nucleotide) and full length NOSIP (912 nucleotide) were synthesized by GenScript and cloned in pET22b expression vector. The recombinant pET22b: iNOS and pET22b: NOSIP plasmid was used to transform competent E. coli BL21 (DE3) cells. The protein was over expressed using 0.2 mM IPTG at 25°C. The harvested cells were resuspended and sonicated in 20 mM Tris-HCl, pH 7.5, 200 mM NaCl, 0.05% Triton X-100 and protease inhibitor (Sigma). The expressed proteins were purified by nickel nitrilotriacetic acid column using 250 mM imidazole.

### Physical interaction by Surface Plasmon Resonance

Interaction of purified NOSIP (full length) with iNOS protein (oxygenase domain) was monitored using Bio-Rad XPR 36 surface Plasmon resonance biosensor instrument. About 3 *µ*g/ml of iNOS was immobilized on GLC chip by amine coupling method as suggested by manufacturer’s instructions. NOSIP was injected at concentration of 500nM with running buffer composed of PBST and 0.005% Tween-20 at a flow rate of 50 *µ*l/min. Molecular interaction was carried out at 20°C. Further kinetic parameter were determined, after fitting the association and dissociation curves to a 1:1 (Langmuir)-binding model. An activated channel without immobilized ligand was used to evaluate nonspecific binding. The response curves were also recorded on control surfaces. Results were calculated after subtraction of the control values using the ProteOn Manager software.

### Isothermal titration calorimetry (ITC)

The iNOS and NOSIP protein were dialyzed in running buffer (reference cell) over night at 4°C and used for protein interaction studies by ITC. The thermodynamics parameters such as Gibb’s free energy change (ΔG), entropy change (ΔS), enthalpy change (ΔH), and the number of binding sites (N) of NOSIP in iNOS binding solution were investigated by isothermal calorimetry MICROCAL PEQ (Malvern) at 25°C. The sample cell was filled with iNOS (10 *µ*M) and the reference cell was filled with 25mM HEPES and 150 mM Sodium chloride, pH=7 and the syringe was filled with NOSIP solution (200 *µ*M). The experiment consisted of 19 injections and each injection contained 1.5 *µ*l of binding solution with 120 second spacing time between subsequent injections. Sequential titrations were carried out and the sample solution was continuously stirred at 270 rpm by the rotating paddle attached to the syringe needle. Multiple injections of NOSIP made into the sample cell containing iNOS. The titration curves were analyzed using the analysis software provided with the calorimeter.

### Modulation of iNOS enzymatic activity by NOSIP

The iNOS recombinant protein (full length) was procured from sigma and Nitric Oxide Synthase activity assay kit was from Abcam. The NOSIP (full length) and iNOS recombinant proteins (full length) were added exogenously into the reaction mixture at 37°C. The enzyme assay was done according to manufacturer’s instructions with addition of Griess reagent. Both iNOS and NOSIP proteins ware equilibrated with 25 mM HEPES, pH=7.4, 150 mM NaCl, 1mM CaCl2 buffer and their concentrations were determined by Bradford assay. 0.5 *µ*M of iNOS was mixed with different concentrations (0.5 to 5 *µ*M) of NOSIP protein and incubated on ice for 30 minutes. The reaction volume was adjusted to 60 *µ*l with NOS assay buffer and another component was added sequentially according to given instructions. The absorbance was measured at 540 nM in microplate reader after reaction completion.

### NOSIP knockdown and functional study

THP-1 cells were grown in complete RPMI medium with 10% FBS. NOSIP shRNA plasmids were obtained from Sigma (MISSION shRNA, cat. No. NM 015953). A cocktail of three different shRNA plasmids was used. Nucleofection was carried out in a Amaxa Nucleofector II machine according to manufacturer’s protocols with slight modifications. Briefly, 5 *µ*g of NOSIP shRNA plasmids or control TRC2 plasmid was mixed with 100 *µ*l Cell Line Nuceleofector Solution V (Lonza) and added to one million cells. For nucleofection, preset program V – 001 was used. Post nucleofection, cells were allowed to grow in complete RPMI media for 48 hours followed by which cells were washed and stimulated with 1 *µ*g/ml LPS for an additional 24 hours. After stimulation, total nitrate + nitrite was estimated in the culture supernatant by Griess assay.

### Identification of NOSIP gene polymorphism

Genomic DNA from whole human blood was isolated using a commercially available kit (Sigma) according to manufacturer’s protocol. The NOSIP gene is present on human chromosome number 19 and spans seven exons. A 503bp segment spanning exon 2 was amplified and tested for novel polymorphism in a cohort of 49 sepsis cases. Primer sequences for amplification are: Forward - 5’ TCCCCATATTCCCACCAGTTTC 3’; and Reverse - 5’ GCCGATGCTAGCTACCACTTGA 3’. PCR was performed for 20 *µ*l reactions containing 2 *µ*l genomic DNA, 1X PCR buffer containing MgCl2 (Sigma), 250 *µ*M dNTP (Sigma), 10 pM of each of forward and reverse primers and 1U of Taq DNA polymerase (Thermo Scientific). PCR cycling conditions were as follows: an initial denaturation step at 94°C for 2 minutes; followed by 35 cycles of 94°C for 15 seconds, annealing at 56°C for 30 seconds and extension at 72°C for 30 seconds; and a final extension at 72°C for 5 minutes. Amplification was checked by running PCR products on a 2% agarose gel. The amplified PCR products were then sequenced on a Genetic Analyzer 3500 DNA sequencing platform (Applied Biosystems).

### RNA extraction and qRT-PCR

Total RNA was extracted from mouse or human whole blood using RNA blood mini kit (Qiagen). Isolated RNA was converted to complementary DNA (cDNA) using RT^2^ first strand cDNA synthesis kit (Qiagen). The cDNA thus obtained was subjected to quantitative real-time PCR analysis with RT^2^ SYBR® Green qPCR Mastermix (Qiagen) in 20 *µ*l reaction volumes on a LightCycler 480 thermal cycler (Roche).

### qRT – PCR array

Human whole blood was stimulated for 4 hours with LPS at 1*µ*g/ml. Following stimulation, total RNA was extracted using a RNA blood mini kit (Qiagen). Total RNA to cDNA conversion was done using RT^2^ first strand cDNA synthesis kit (Qiagen). Prepared cDNA was subjected to a customised qRT – PCR array (SABiosciences) using RT^2^ SYBR® Green qPCR Mastermix (Qiagen). Fold change over untreated control was calculated using the 2^-ΔΔCt^ method. The obtained fold change values were used for pathway enrichment analysis (Ingenuity Systems).

### Statistical analysis

For comparison between two groups, either an unpaired t – test or the nonparametric Man – Whitney U –test was conducted. One – way analysis of variance (ANOVA) followed by Bonferroni’s multiple comparison test was carried out to compare means between three or more groups. Difference in survival among different groups was estimated by constructing Kaplan – Meier survival curves followed by assessment of significant difference using the log – rank test. Fisher’s exact test was used for comparison of genotype, allele frequencies and to test association of combined genotype distribution among various clinical categories. For all statistical comparisons, a p-value *<*0.05 was considered statistically significant.

## Results

### NOSIP interacts with iNOS and modulates nitric oxide synthesis

Since binding of NOSIP to eNOS (endothelial NOS, NOS3) and nNOS (neuronal NOS, NOS1) and its inhibition of their function has been reported earlier, the first objective of the current study was to investigate whether NOSIP interacts with iNOS. To test for direct physical interaction between the two molecules in a cell free system, we performed surface plasmon resonance (SPR) studies with recombinant iNOS and NOSIP. Our results demonstrate a K_D_ value of 7.11*X*10^*−*9^*M* that is indicative of a significant interaction between the two proteins (Figure 1A).

**Figure 1.**
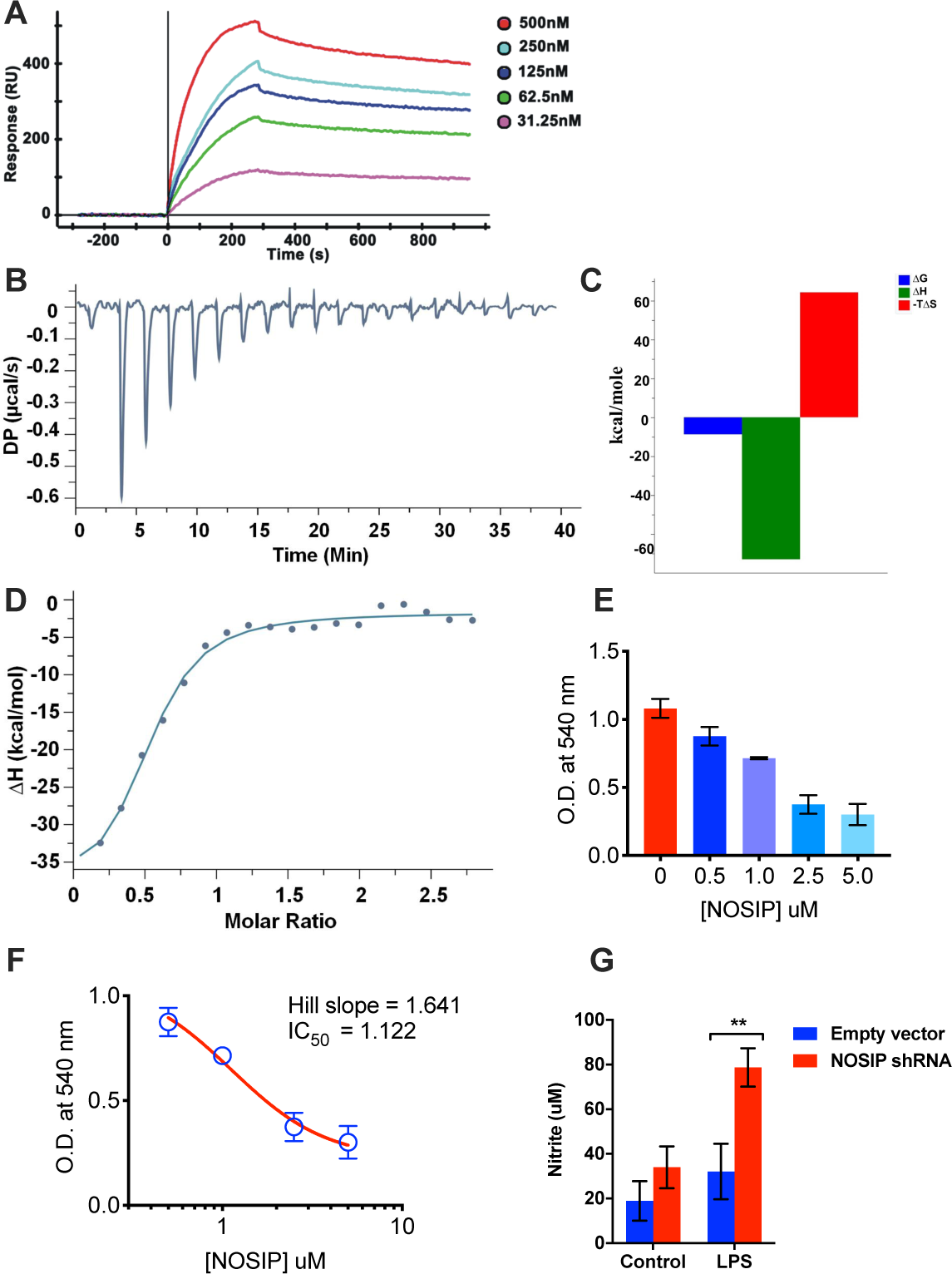
NOSIP (full length) interacts with iNOS (oxygenase domain) and regulates nitric oxide synthesis. (A) Dose dependent binding of NOSIP recombinant protein (analyte) to immobilized iNOS solid phase (ligand) was measured by surface plasmon resonance. Recombinant NOSIP protein was injected at concentrations of 500nM, 250nM, 125nM, 62.5nM to 31.25nM over ligand. The *K*_D_ = 7.11*e−* 09*M* was obtained in Kinetic – Langmuir analysis. (B) Calorimetric titration of NOSIP (full length) and iNOS (Oxygenase domain) by ITC: heat change per injection of NOSIP to iNOS are shown. (C) Graphical representation of thermodynamic parameters from the titration experiments in B. (D) The resulting isotherms from C fitted with one site binding model. *In vitro* iNOS enzymatic activity was tested individually and in combination with NOSIP. Red bar - iNOS enzyme activity in the absence of NOSIP, measured by nitrite estimation from substrate in a reaction. The different blue bars show titration of different concentrations of NOSIP inhibiting iNOS enzymatic activity. (F) Dose response curve fitting of NOSIP mediated inhibition of iNOS. The data from (E) was fitted using a four-parameter Hill function. (G) THP-1 cells were nucleofected with empty plasmid or NOSIP shRNA plasmid by Amaxa nucleofector II using manufacturer’s kit. After 48 hours, the cells were washed and incubated with or without LPS at 1 *µ*g/ml for a further 24 hours. Total nitrate+nitrite in culture supernatant was measured by a commercial Griess assay kit. Statistical significance was assessed by two – way ANOVA following Bonferroni’s post test (** p*<*0.01). The graph represents data from 5 separate plates performed on 2 different days.

Further confirmation of direct binding between iNOS and NOSIP came from ITC titration profile of NOSIP binding with iNOS as shown in Figure 1B-D. The thermodynamic parameters such as entropy change (-TΔS) was found to be 29.9 kcal/mol, enthalpy change (ΔH) −38.20 kcal/mol, Gibb’s free energy change (ΔG) was −8.3 kcal/mol and the number of binding sites (N) of NOSIP in iNOS binding solutions was around 0.505. Reaction of iNOS with NOSIP was exothermic in nature as suggested by the negative value of enthalpy change. The next critical objective was to test biochemical consequence of interaction between iNOS and NOSIP on synthesis and release of Nitric Oxide. We measured NOSIP mediated inhibition of iNOS in a cell-free system. We found NOSIP diminished the enzymatic activity of iNOS in a dose-dependent manner (Figure 1E) with an IC50 value of 1.12 *µ*M, as calculated by fitting the data to a four-parameter Hill function (Figure 1F). Next, to validate our results in a cell-free system and to check for functional consequence of iNOS-NOSIP interaction, we performed targeted knockdown of NOSIP in THP-1 human monocytic cell line. Stimulation of THP-1 cell lines with LPS for 24 hours revealed significantly higher levels of nitrite in culture supernatants of cells transfected with NOSIP shRNA as compared to cells transfected with null vector (Figure 1G), suggesting that NOSIP inhibits nitric oxide production likely through its interaction with iNOS. Confocal imaging analysis of iNOS and NOSIP proteins showed co–localization of the proteins in the human monocytic cell line THP-1 and primary human monocytes demonstrating similar subcellular localization (Figure S1A and B). In situ proximity ligation analysis of iNOS and NOSIP in primary human monocytes further confirmed a direct physical interaction between iNOS and NOSIP (Figure S1C). Interestingly, we observe considerable nuclear localization of both iNOS and NOSIP (Figure S1), which could be a potential mechanism by which NOSIP inhibits iNOS.

Taken together, above findings demonstrate that NOSIP interacts with iNOS and modulates synthesis of nitric oxide.

### A four bp deletion downstream of Exon 2 in NOSIP gene leads to increased expression of NOSIP protein in human peripheral blood cells

Since NOSIP regulates NO production we searched for possible polymorphisms in NOSIP gene that may have functional consequences in inflammatory responses in humans. Five different sets of primers each amplifying different regions of nosip gene were designed and the amplified products were sequenced (Figure S2). Sequence matching of 59 DNA samples isolated from healthy volunteers with that of the database revealed presence of a four nucleotide deletion (TCTC) downstream of exon two in about 71% (42/59) of the subjects (Table 1). Next we addressed the issue whether the polymorphism influences NOSIP expression - intracellular NOSIP levels between ‘insertion’ and ‘deletion’ individuals was scored in circulating leucocytes of healthy human volunteers by imaging cytometry (Figure 2). In the first plot, a histogram of brightfield channel intensity is used to exclude ‘out-of-focus’ cells by visually inspecting each bin. The resulting ‘properly focused’ population (R1) is then plotted for brightfield area and aspect ratio to include only single cells (R2) and exclude doublets (high area) and debris (low area and low aspect ratio). Subsequently, a bivariate scatterplot of the singlet population based on side scatter profile and intracellular NOSIP expression revealed three distinct cell populations – regions R7, R8 and R9 (Figure 2A). Figure 2B-D shows representative brightfield, NOSIP and nuclear images of cells drawn from each of R7, R8 and R9. Cells within the R9 gate displayed the highest NOSIP expression. Comparison of intracellular NOSIP expression in all three populations between ins and del individuals revealed increased expression in individuals with deletion polymorphism as compared to those with insertion (Figure 2E-G). In summary, above data provide evidence for existence of a novel polymorphism in NOSIP gene that leads to higher intracellular expression of NOSIP in circulating immune cells.

**Table 1.** Genotype and allele distribution of NOSIP (TCTC ins/del) polymorphism in healthy individuals and patients with sepsis.

| Genotype | Healthy controls (n = 59) | Sepsis (n = 45) |  |
| --- | --- | --- | --- |
| *ins/ins* | 17 (28.2%) | 19 (42.2%) |  |
| *ins/del* | 28 (47.5%) | 18 (40.0%) |  |
| *del/del* | 14 (23.7%) | 8 (17.8%) |  |
| Allele | Healthy controls (n = 118) | Sepsis (n = 90) |  |
| *ins* | 62 (52.5%) | 56 (62.2%) |  |
| *del* | 56 (47.5%) | 34 (37.8%) |  |
| Genotype | P-value | Odds ratio | 95% C.I. |
| *ins/ins* vs. *ins/del* | 0.2662 | 0.5752 | 0.2512 to 1.416 |
| *ins/ins* vs. *del/del* | 0.2828 | 0.5113 | 0.1864 to 1.441 |
| Allele | P-value | Odds ratio | 95% C.I. |
| *ins* vs. *del* | 0.2036 | 0.6722 | 0.3920 to 1.177 |

**Note:** The percentages in parentheses denote the overall distribution of a particular genotype or allele in a given category. The odds ratio is calculated by a Fisher’s exact test to compare the distributions of a category of patients between two genotypes or alleles. A p-value <0.05 is considered statistically significant.

**Figure 2.**
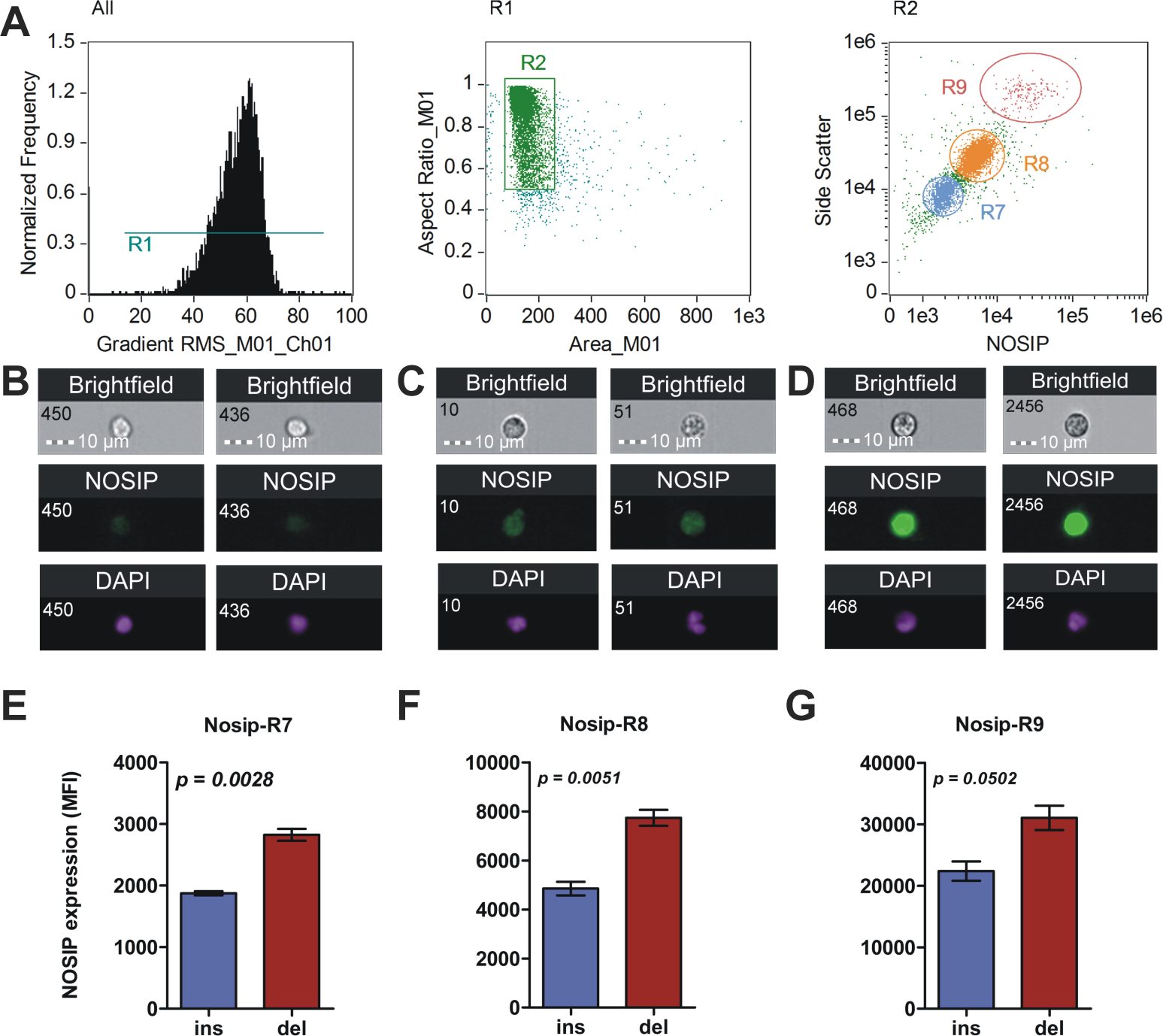
A novel mutation in NOSIP gene leads to increased intracellular protein expression. (A) Imaging cytometry analysis of intracellular NOSIP expression. In the first plot, the histogram depicts selective removal of ‘out – of – focus’ cells. In the next plot, only single cells are selected for further analysis (R2). Finally, they were discriminated on a bivariate scatter plot based on differences in side scatter signal and intracellular NOSIP expression (regions R7, R8 and R9). (B – D) Representative imgaes showing cell and nuclear morphology along with intracellular NOSIP expression of populations R7 (B), R8 (C) and R9 (D). (E – G) Graphs showing significantly elevated levels of NOSIP protein as a consequence of presence of deletion allele as assessed by unpaired t – test (n = 4 for ins and n = 3 for del category). Statistical significance was assessed by an unpaired t-test with Welch’s correction.

### Higher NOSIP expression in humans is associated with increased inflammation and susceptibility to acute inflammatory diseases

We then wanted to test the functional significance of NOSIP mutation in host inflammatory responses. Whole blood of normal human volunteers were stimulated with LPS for 4 hours and gene expression was compared between ins or del subjects by a custom designed qRT-PCR array (Figure 3A) – very broadly, expression of many of the cytokine and chemokine genes in subjects with del/del genotype were higher in comparison to those with Ins/Ins genotype (Cluster 4, Figure 3A). In order to gain insights into global differences in signalling pathways as a consequence of differential NOSIP expression, a pathway enrichment analysis was performed. The analysis revealed increased enrichment of pathways associated with inflammatory response, viz. Toll-like receptor signalling, NF-*κ*B signalling, p38 MAP kinase signalling and HMGB1 signalling in individuals with del allele (Figure 3B). A similar enrichment analysis conducted for disease associated networks revealed higher enrichment of sepsis, septic shock network in Del individuals. Higher representation of networks associated with liver and kidney damage, apoptosis of liver and kidney cells, and apoptosis of macrophages, phagocytes and antigen presenting cells were also observed suggesting an increased risk of acute inflammation associated pathology in individuals with increased expression of NOSIP (Figure 3C). Visual comparison of septic shock associated networks between Ins/Ins (Figure 3D) and Del/Del (Figure 3E) individuals confirmed this. Therefore, an analysis of NOSIP polymorphic genotype distribution was performed in a cohort of patients with sepsis. Although there was no significant difference in distribution of NOSIP mutation between controls and sepsis patients (Table 1), the results revealed significantly higher mortality among sepsis patients displaying homozygous deletion polymorphism. Significantly high odds ratio (5/8 deaths, P-value = 0.0267, OR = 8.889, 95% C.I. = 1.344 to 58.8) indicated that patients with homozygous deletion mutation are highly prone to mortality in the cohort of sepsis patients studied (Table 2).

**Table 2.** Genotype and allele distribution of NOSIP (TCTC ins/del) polymorphism and its association with mortality in sepsis.

| Genotype | Survivors (n = 32) | Nonsurvivors (n = 13) |  |
| --- | --- | --- | --- |
| *ins/ins* | 16 (50.0%) | 3 (23.1%) |  |
| *ins/del* | 13 (40.6%) | 5 (38.5%) |  |
| *del/del* | 3 (9.4%) | 5 (38.5%) |  |
| Allele | Survivors (n = 64) | Nonsurvivors (n = 26) |  |
| *ins* | 45 (70.3%) | 11 (42.3%) |  |
| *del* | 19 (29.7%) | 15 (57.7%) |  |
| Genotype | P-value | Odds ratio | 95% C.I. |
| *ins/ins* vs. *ins/del* | 0.4470 | 2.051 | 0.411 to 10.24 |
| *ins/ins* vs. *del/del* | **0.0267** | 8.889 | 1.344 to 58.8 |
| Allele | P-value | Odds ratio | 95% C.I. |
| *ins* vs. *del* | **0.0171** | 3.23 | 1.255 to 8.309 |

**Note:** The percentages in parentheses denote the overall distribution of a particular genotype or allele in a given category. The odds ratio is calculated by a Fisher’s exact test to compare the distributions of a category of patients between two genotypes or alleles. A p-value <0.05 is considered statistically significant.

**Figure 3.**
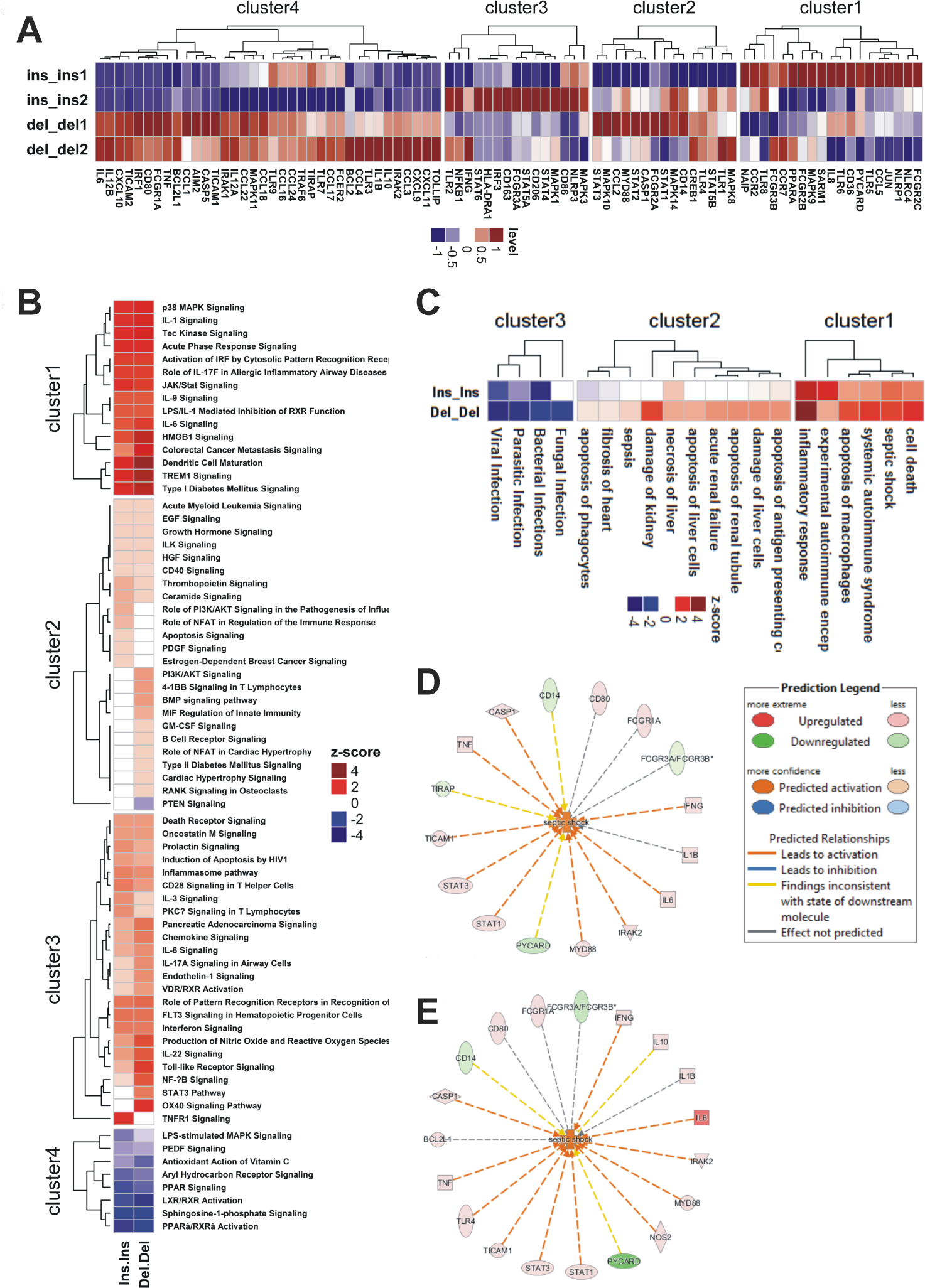
Comparison of global changes in gene expression and pathways between Ins and Del individuals. Whole blood was collected from two healthy individuals each of ins/ins and del/del genotypes and stimulated with LPS at 1 *µ*g/ml for 4 hours. After stimulation, total RNA was extracted, cDNA synthesized and subjected to a custom designed RT2 q-PCR array. Raw Cp values were exported in Qiagen’s online tool and fold increase was calculated after normalizing with *β*-actin as housekeeping gene. **(A)** Clustering heatmap depicting differential expression of 85 immune response genes between ins/ins and del/del individuals. **(B)** Differences in pathway enrichment between ins/ins and del/del individuals as assessed by Ingenuity Pathway Analysis. **(C)** Differential enrichment of disease – associated networks between ins/ins and del/del individuals. (**D and E)** Representative networks associated with septic shock response between ins/ins **(D)** and del/del **(E)** individuals.

Taken together, the above results clearly demonstrate that individuals homozygous for the deletion allele display increased inflammatory features and are more susceptible to acute inflammation induced morbidity and mortality.

### NOSIP levels are associated with differential response to LPS activation between circulating monocytes of mouse and human origin

There is evidence in literature about a species-specific hierarchy in susceptibility to acute inflammation [37]. Our next objective was to test whether NOSIP levels are associated with species-specific variability in a host’s response to inflammatory insult. A comparison of intracellular iNOS and NOSIP expression between circulating monocytes of human and mouse revealed significantly higher levels of iNOS (Figure 4A) and NOSIP in human monocytes (Figure 4B). In concordance with the above data, we also observed significantly lower levels of nitrite in plasma of sepsis pateints as compared to plasma of mice injected with LPS (Figure S3). We then wanted to test the functional consequence of differential expression of NOSIP between mouse and human immune cells upon inflammatory activation. As such, human and mouse whole blood were stimulated with LPS and intracellular IL-1*β* and TNF-*α* were measured by flow cytometry. For the purpose of comparison, we tested CD14(+)/Ly-6G(-) cells in mice blood with that of CD14dim/C16hi nonclassical monocytes in humans, demonstrated to be the most activated monocyte subtype in humans [38]. Our results show that human peripheral blood monocytes produce significantly higher IL-1*β* as compared to monocytes of mice upon stimulation with LPS (Figure 4C and 4E). Interestingly, however, TNF-*α* levels were comparable between the two species as depicted in Figure 4D and 4E.

**Figure 4.**
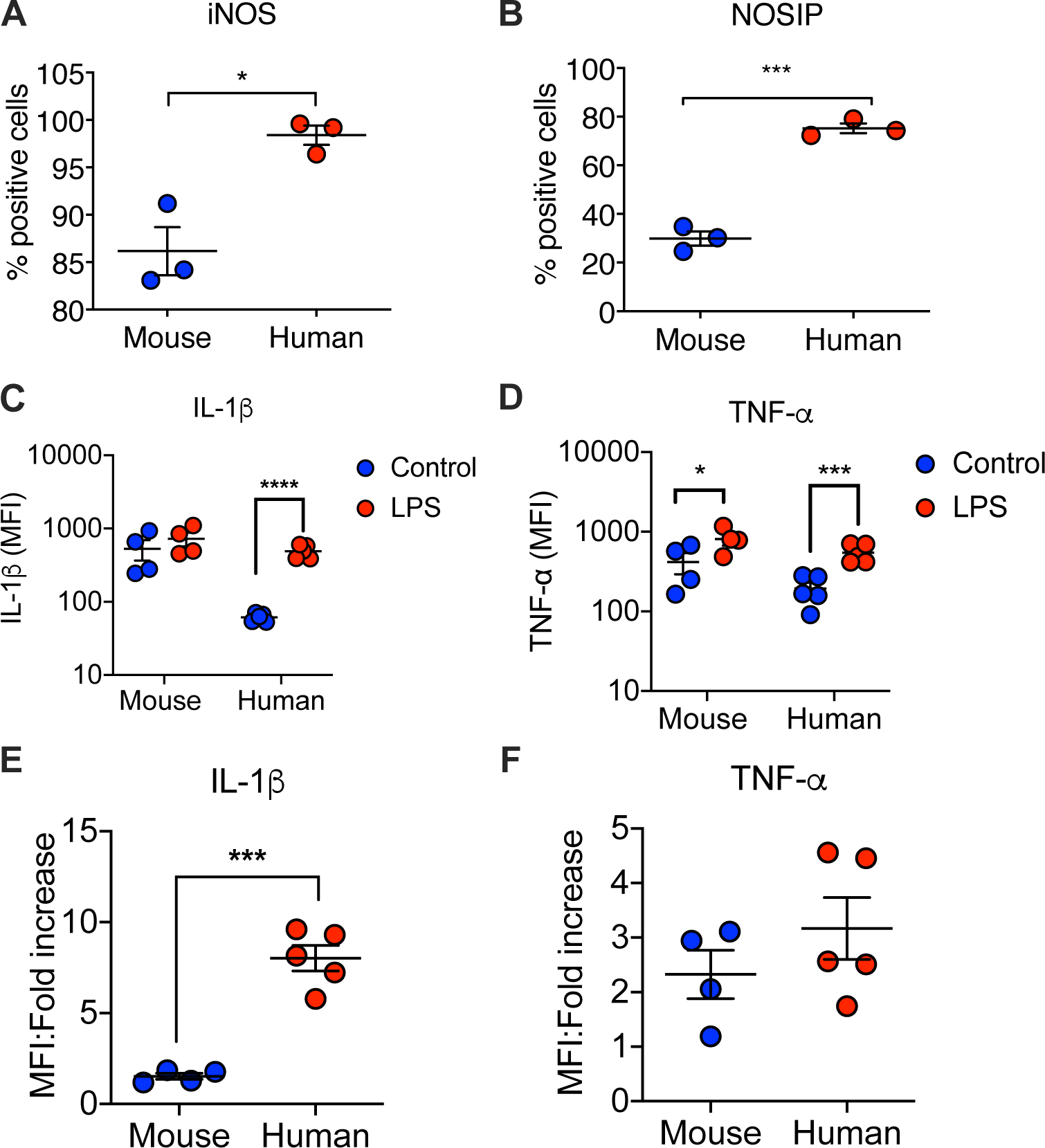
Species-specific differences in NOSIP expression regulates response to LPS. (A) Comparison of intracellular protein expression of iNOS in human and mouse circulating CD14+ monocytes. (B) Comparison of intracellular protein expression of NOSIP in human and mouse circulating CD14+ monocytes. For both A and B, n = 3 for both mouse and human. (C - F) Mouse (n = 4) and human (n = 5) whole blood were left untreated or treated with 1 *µ*g/ml LPS for 2 hours along with Brefeldin A. Following stimulation, cells were fixed, permeabilized, washed and stained with flurochrome-conjugated antibodies for analysis on a flow cytometer. (C and D) Comparison of LPS-induced IL-1*β* and TNF-*α* between mouse and human monocytes. (E and F) The data in (C) and (D) depicted as fold change over untreated control. (C - F) For mouse, cells gated on CD14(+)/Ly-6G(-) were used for analysis; for humans, we used a CD14dim/CD16hi nonclassical monocyte subset. (A, B, E, F) Statistical significance was tested using an unpaired t-test. (C and D) Statistical significance assesed by multiple comparison t-test with Holm-Sidak correction. * *p<0*.*05*, \*\*\**p<0*.*001*, \*\*\*\**p<0*.*0001*.

A key point to note here is that inbred SPF mice are very different in terms of their physiology from humans due to subclinical infections or vaccinations. Moreover, experimentally induced endotoxemia in mice is not similar to human sepsis. However, in spite of the above caveats, the above observations indicate that differential synthesis of nitric oxide (as a consequence of differences in NOSIP expression) between species may be in part responsible for regulating the response to acute inflammatory stimulus.

### Plasma nitrite levels are inversely associated with inflammatory cytokines in human sepsis and experimental murine endotoxemia

The observation that increased NOSIP leads to decreased NO synthesis may be harmful to host during acute inflammation led to us to investigate the role played by NO during acute inflammatory disorders. In order to gain insight into the role of nitric oxide in acute inflammation, a multivariate correlation analysis of plasma cytokines and nitrite levels was conducted in human sepsis patients. A significant negative association was observed between plasma nitrite and some of the cytokines/chemokines (Table 3) suggesting a possible protective role for increased plasma nitrate/nitrite levels in human sepsis. To test this hypothesis experimentally, male C57BL/6 mice were injected with two different doses of LPS and plasma cytokines and nitrite levels were estimated at 2, 4, 8 and 16 hours post injection. Plasma nitrite levels negatively correlated with inflammatory mediators in mouse model of experimental endotoxemia (Table 3).

**Table 3.** Inverse association of plasma nitrite with inflammatory factors in a mouse model of endotoxemia and human sepsis.

| Factor | By factor | Human |  | Mouse |  |
| --- | --- | --- | --- | --- | --- |
| | | Spearman $\rho$ | p-value | Spearman $\rho$ | p-value |
| Nitrite | IL-1β | -0.17808 | 0.03805 | -0.63321 | <0.0001 |
| Nitrite | IL-6 | -0.06230 | n.s. | -0.29015 | 0.03006 |
| Nitrite | IL-8 | -0.18577 | 0.03036 | N.A. | N.A. |
| Nitrite | IL-9 | -0.16614 | 0.05322 | N.A. | N.A. |
| Nitrite | IL-12 (p70) | 0.08337 | n.s. | -0.32543 | 0.01438 |
| Nitrite | MIP-α | -0.25125 | 0.00317 | -0.62772 | <0.0001 |
| Nitrite | MIP-1β | -0.14636 | n.s. | -0.61086 | <0.0001 |
| Nitrite | TNF-α | -0.10051 | n.s. | -0.72785 | <0.0001 |

**Note:** Plasma nitrate+nitrite was estimated by a commercially available kit. Cytokines in plasma was measured by Bioplex suspension array system. Only cytokines having negative association with nitrite are shown. The correlation values displayed were calculated by constructing a $n \times n$ scatterplot matrix and scoring for bivariate associations between each pair of factors. *N.A.* Not available; *n.s.* not significant.

### Deficiency in nitric oxide synthesis increases susceptibility to acute inflammation

From the above data, we hypothesised that absence of nitric oxide could exacerbate an inflammatory insult. To investigate this, male BALB/C mice were treated with 100 mg/kg of N*ω*-Nitro-L-arginine methyl ester (L-NAME) hydrochloride once every 24 hours to block nitric oxide synthase activity. L-NAME is a structural analogue of L-Arginine that competitively inhibits all three isoforms of nitric oxide synthase. Untreated and L-NAME treated mice were then injected with 2 mg/kg of LPS and mortality was assessed for 96 hours. Figure 5A shows comparison in LPS toxicity between untreated and L-NAME treated animals clearly demonstrating that inhibition of nitric oxide synthesis is detrimental to host in a mouse model of endotoxemia. This was further tested by studying LPS toxicity in mice deficient in Nos2 gene. Comparison of lethal dose of LPS between wild type C57BL/6 mice and Nos2 null revealed that 15mg/kg of LPS did not result in mortality of wild type mice, while about 84% of the animals in the Nos2−/− group died at the same dose of LPS (Figure 5B). This confirmed the hypothesis that absence of nitric oxide inducing enzyme and consequently nitric oxide results in increased susceptibility to endotoxemia. To test for possible mechanisms of the observed phenomenon, wild type and Nos2−/− mice were injected with LPS at a dose of 15mg/kg and the animals were sacrificed at 2, 6 and 12 hours post administration and plasma cytokines were quantified. The heatmap shown in Figure 5C reveals significantly higher levels of IL-1*β* in mice lacking the Nos2 gene. However, plasma levels of TNF-*α* were comparable in both (Figure 5C), suggesting that IL-1*β* could be playing a more central role in poor disease outcome in endotoxemia/sepsis and that NO is a key molecule regulating NLRP3 mediated inflammasome pathway. Very similar observations on the role of nitric oxide on IL-1*β* in endotoxemia has been reported earlier [34]. In order to obtain a more global understanding of the effect of Nos2 deficiency in LPS induced inflammation, we performed a principal component analysis (PCA) of the measured cytokines. The scatterplot depicted in Figure 5D clearly shows the best separation of wild type and Nos2−/− along the third principal component. This is also evident from the scores on principal component 3 (PC3), which shows a clear dichotomy between wild type and Nos2−/− mice (Figure 5E). Analysis of the contribution of individual cytokines along PC3 revealed IL-1*β* to be the highest positive contributor towards the observed differences between wild type and Nos2−/− mice (Figure 5F). Moreover, we also observed significantly higher levels of IL-5 in Nos2−/− mice along with a concordant decrease in IL-12 and IFN-*γ* levels (Figure 5C and F). This indicates a possibility of a T-helper type2 skewed response in the Nos2−/− mice.

**Figure 5.**
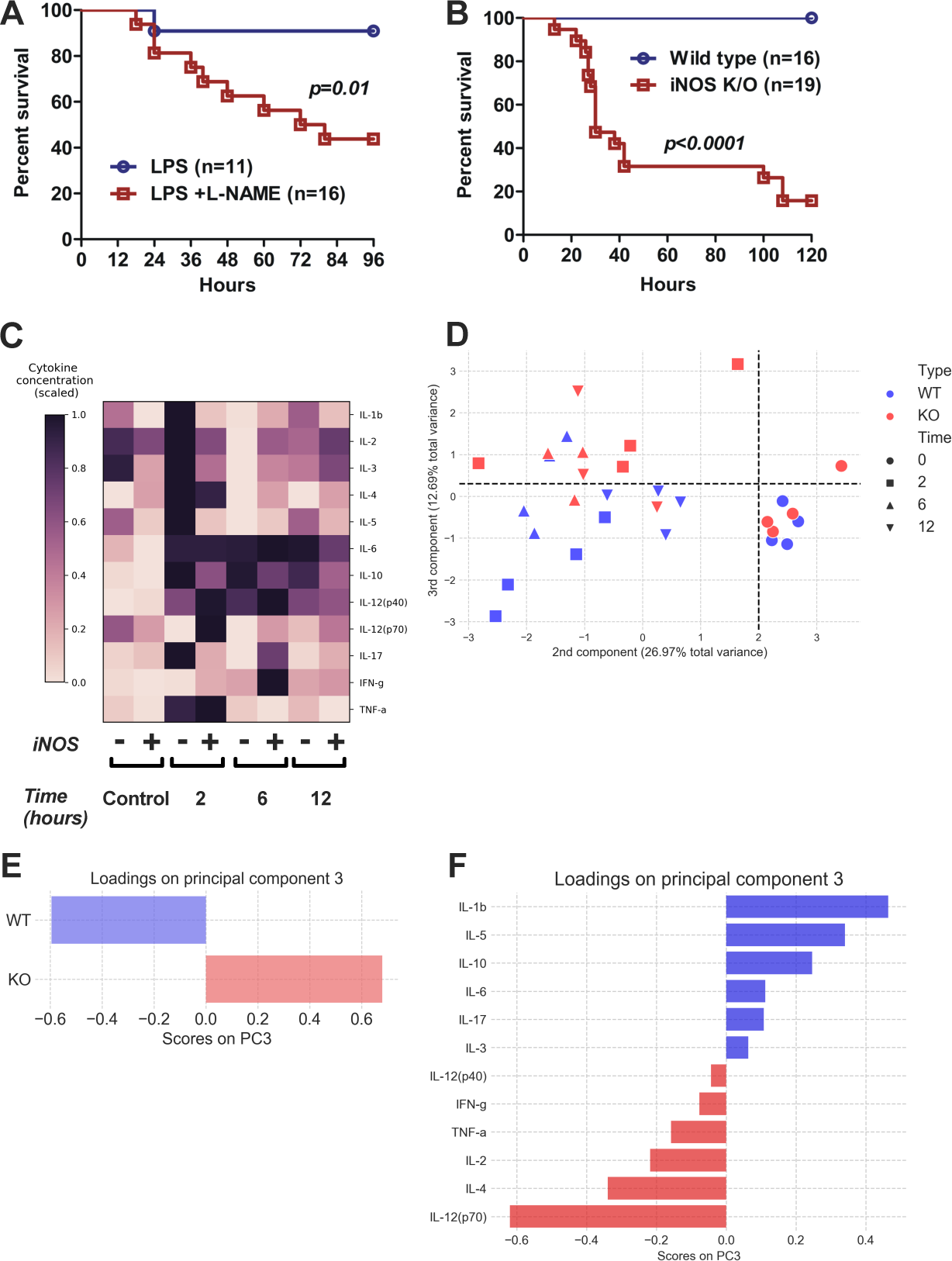
Deficiency of nitric oxide increases LPS – mediated toxicity involving increased IL-1*β* secretion. **(A)** Treatment of BALB/C mice with a nonselective NOS inhibitor renders the animals more susceptible to LPS – mediated endotoxic shock in comparison to untreated mice. L-NAME was injected at 100 mg/kg once every 24 hours. LPS was injected at 2 mg/kg body weight. **(B)** iNOS−/− C57BL/6 mice and their wild-type littermates were injected with 15 mg/kg LPS and mortality was scored for 120 hours. **(C)** Heatmap depicting the time kinetics of plasma cytokines in LPS treated W/T and iNOS−/− mice. The animals (n = 4 in each group) were weighed and LPS was injected at 15 mg/kg intraperitoneally. For assessment of cytokine production, animals were sacrificed at the indicated time points and isolated plasma was subjected to a multiplex cytokine assay. The x-axis labels show the time pint followed by the genotype. Each voxel in the heatmap is a mean of 4 mice. The values are scaled to reduce the dynamic range in colouring scheme. Significance levels were assessed by two-way ANOVA with Bonferroni’s multiple comparison test. **(D)** Scatterplot showing the distribution of the samples in a space marked by principal components 2 and 3 from a principal component analysis of plasma cytokines in LPS injected W/T and iNOS−/− mice. **(E)** Barplot showing difference in score for PC3 for the two genotypes tested. Each bar is a combination data from all time points and is a mean of n = 12 mice. **(F)** Factor loadings plot showing the contribution of individual cytokines towards PC3.

To summarize, the above findings demonstrate that absence of Nos2 is deleterious to host in a mouse model of acute inflammation that involves increased production of IL-1*β*. Figure 6 summarizes the principal findings of this study.

**Figure 6.**
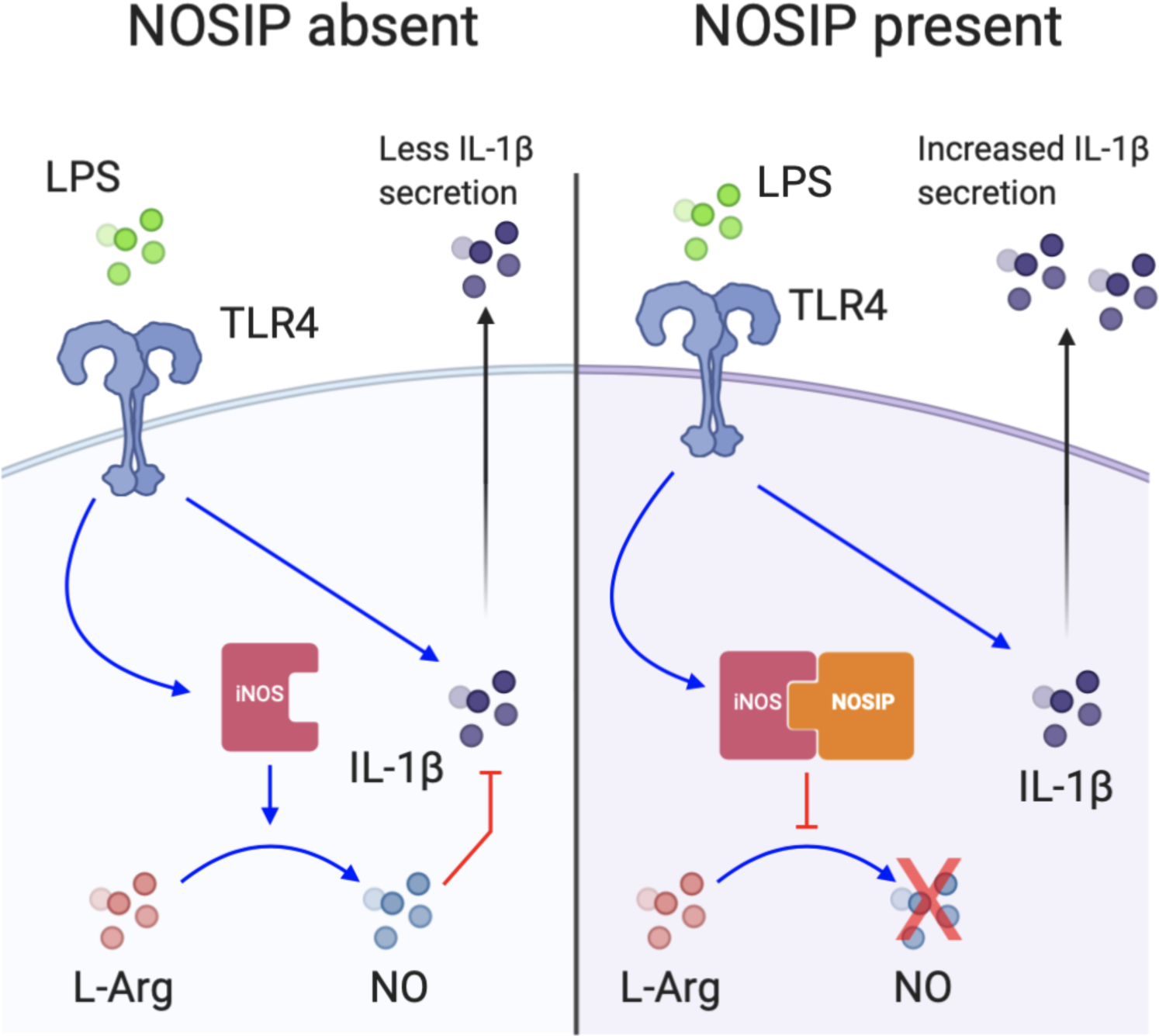
Model of NOSIP-mediated regulation of LPS response. Ligation of LPS-CD14 complex (not shown for sake of simplicity) to membrane TLR4, leads to activation of iNOS along with IL-1*β*, that is secreted from the cell. *(Left)* In the absence of NOSIP protein, iNOS is available to catalyze the conversion of L-Arginine to nitric oxide (NO), which then inhibits IL-1*β* production, thereby leading to less IL-1*β* secretion. *(Right)* When NOSIP is present, it binds to iNOS and inhibits its ability to convert L-Arginine to NO. Less nitric oxide leads to increased production and secretion of IL-1*β*.

## Discussion

NOSIP has diverse biological functions. Originally discovered as a protein interacting with e-NOS in endothelial cells and regulating the enzyme in its translocation from plasma membrane leading to decreased production of Nitric Oxide [27], it was subsequently demonstrated to regulate neuronal functions by interacting with nNOS [26]. More recently it has been shown to be a ubiquitin E3 ligase modulating protein phosphatase 2A and playing a role in development of vertbrate eye and cranial cartilage has been proposed [39, 40]. Curiously however the role of NOSIP in regulating iNOS and its consequence in mammalian immune system has not been investigated. We provide unequivocal biochemical and biophysical evidence (apart from in situ microcopic and functional proof) for interaction between iNOS and NOSIP using a cell free system using recombinant protiens, unlike previous investigations on eNOS/nNOS and NOSIP, that mostly utilized yeast two-hybrid screens and chromatin immunoprecipitation assays [26, 27].

Nitric oxide (NO) is synthesised by many cell types during host response to pathogens and injury [4]. The principal enzyme responsible for nitric oxide production in immune cells during inflammation is inducible nitric oxide synthase (iNOS or NOS2). Role of nitric oxide in endotoxemia/sepsis has been very contradictory in reported literature. Earlier studies suggested a detrimental role for nitroic oxide and as a consequence inhibitors of iNOS were tested for treatment of endotoxemia/sepsis. Although pre-clinical trials offered promising results a human trial conducted with iNOS inhibitor, aminoguanidine had to be truncated since prognosis was better in patients treated with placebo [31]. Viewed from the observations of the current study which offers evidence for an anti-inflammatory role for Nitric Oxide, failure of clinical trial with iNOS inhibitor is not surprising. A recent study further adds credence to protective role of Nitric oxide in sepsis. Therapeutic potency of transfer of ex vivo expanded fibroblastic reticular cells (FRCs) into mice with endotoxemia or cecal-ligation-and-puncture induced sepsis, was found to be dependent on iNOS activity expressed by FRCs [41]. In a similar sepsis model, iNOS-dependent upregulation of cGMP and subsequent activation of TACE was found to protect against organ injury [42]. Other studies in animal models have also documented possible protective role of NO in endotoxemia [43].

The current manuscript presents several novel findings on modulation of iNOS activity and nitric oxide biology in the context of host response by Nitric oxide synthase interacting protein (NOSIP). Several years ago NOSIP was discovered to interact with eNOS and inhibit its function by sequestering it in Golgi complex [27]. Subsequently, interaction of NOSIP with nNOS was also reported [26]. Curiously however, till date the role of NOSIP and its with iNOS and regulation of nitric oxide production by immune cells has not been investigated – the present study fills this lacuna. Physical interaction between NOSIP with iNOS were demonstrable in immune cells as well as in cell free systems using recombinant proteins. More significantly, NOSIP emerged as a key molecule regulating iNOS function and release of nitric oxide. More critically a four nucleotide deletion upstream of first exon of NOSIP gene was found to be associated with increased intracellular expression of NOSIP and higher risk of mortality due to sepsis – clinical severity in which have been associated with enhanced host inflammation. These observations have opened up an entirely new area of therapeutic strategy for management of acute inflammation. Structural biology of iNOS-NOSIP interaction can be expected to lead to discovery of small molecules for blocking their interactions which could modulate nitric oxide levels and consequently inflammation. More significantly such small molecules could also be useful in regulating nitric oxide production by nNOS and eNOS in neuronal and endothelial cells respectively.

An inverse association of plasma nitrite with inflammatory cytokines was observed in human sepsis and mouse model of endotoxemia, suggesting a reciprocal regulation of nitric oxide and inflammatory mediators. Species-specific differences in nitric oxide synthesis have been well documented in literature. A major issue of debate in literature has been differences in NO production among species, most notably between human and mouse [44, 45]. It has been reported earlier that human monocytes and macro-phages produce significantly lower amounts of NO than their mouse counterparts [46]. Whether such differences translate into differences in susceptibility to acute inflammation remains unclear and is one of the key questions addressed in the present study. The results show a clear positive association between nitrite oxide production and resistance to LPS mediated inflammation, suggesting that a given species’ ability to tolerate inflammatory insult is, at least in part, dictated by its ability to produce nitric oxide. This conclusion may have important bearing on a study published earlier that demonstrated that proteins in serum rather than intrinsic cellular differences may play a role in regulating variations in resilience to microbe-associated molecular patterns between species [37]. It is apparent that identification of such proteins may open up new areas of targeted therapy. However, given the large number of proteins in serum, this is an arduous task. The observations made in the current study could help in narrowing down the search by initially probing enzymes involved in nitric oxide biosynthesis.

Another important finding of the present investigation was demonstration of increased IL-1*β* production in mice lacking a functional iNOS gene (during inflammatory activation), suggesting that protective effect of nitric oxide in endotoxemia is mediated through inhibition of IL-1*β* synthesis – more so since TNF-*α* levels remained comparable in wild type and iNOS deficient mice. This is particularly interesting in the light of an earlier study that demonstrated no difference in LPS toxicity between wild type and TNF-*α* knockout mouse [47]. On the other hand, mice deficient in IL-1*β* converting enzyme were relatively resistant in comparison to wild type mice when challenged intraperitoneally with endotoxin [48]. These observations suggest a central role played by IL-1*β* (inflammasome pathway) in pathogenesis in endotoxemia/sepsis. Similar to nitric oxide, species specific dichotomy in terms of IL-1*β* production was observed between human and mouse circulating immune cells that negatively correlated with nitric oxide production, further demonstrating IL-1*β* as a central pathogenic hub in endotoxemia.

Despite advances in both understanding of basic biology as well as better critical care support, mortality due to sepsis remains unacceptably high. As of now, no effective therapy exists to combat sepsis. Given the protective role played by nitric oxide in sepsis, it is tempting to suggest administration of nitric oxide donors for clinical management of sepsis. Indeed, several studies have attempted NO supplementation in sepsis, and a systematic review and meta-analysis of such studies showed that this line of therapy could be promising [49]. However, because of the extremely short half-life of nitric oxide, NO donors require administration on a regular basis, thus potentially raising the cost of treatment. In this context, identification of NOSIP as a key regulator of NO synthesis immediately opens up exciting possibilities for developing targeted therapy. Designing inhibitors of NOSIP could lead to increased NO production which, in turn, could control hyper-inflammation observed in sepsis. Moreover, since nitric oxide acts as a potent vasodilator and an inhibitor of platelet aggregation, the implications of developing inhibitors against a protein that modulates its synthesis are quite attractive as a therapeutic strategy in cardiovascular diseases. However, owing to the novelty of NOSIP, extreme caution should be exercised before designing such therapies as the overall function of NOSIP in other cellular processes need to be evaluated first, particularly their effect on eNOS and nNOS mediated biological activities - a recent finding reported a crucial requirement of NOSIP during neurogenesis in both mice and Xenopus [50]. Since increased NOSIP expression appears to render a host more susceptible to acute inflammation from an evolutionary perspective, it would be interesting to investigate differences in organ specific NOSIP expression between species and strains of animals.

Finally, from a translational perspective the role of this novel NOSIP polymorphism viz., (subjects with homozygous ‘del’ allele being more prone for inflammation), in determining disease severity as well as mortality of the ongoing Covid 19 pandemic will be of immediate relevance to be used as a biomarker in Covid 19 infections – host inflammation is a key factor in pathogenesis of Covid 19 infections [51] and use of anti-inflammatory drugs for clinical management of Covid 19 have been proposed [52].

## Acknowledgments

The authors thank Dr. Dileep Vasudevan, Institute of Life Sciences for his assistance in cloning and purification of iNOS and NOSIP proteins and Dr. Narottam Acharya, Institute of Life Sciences for his assistance in interaction studies by Surface Plasmon Resonance. The Institute of Life Sciences is funded by grants from Department of Biotechnology, Govt. of India. R.M. is a recipient of senior research fellowship from Indian Council of Medical Research.

---

**Figure S1.**
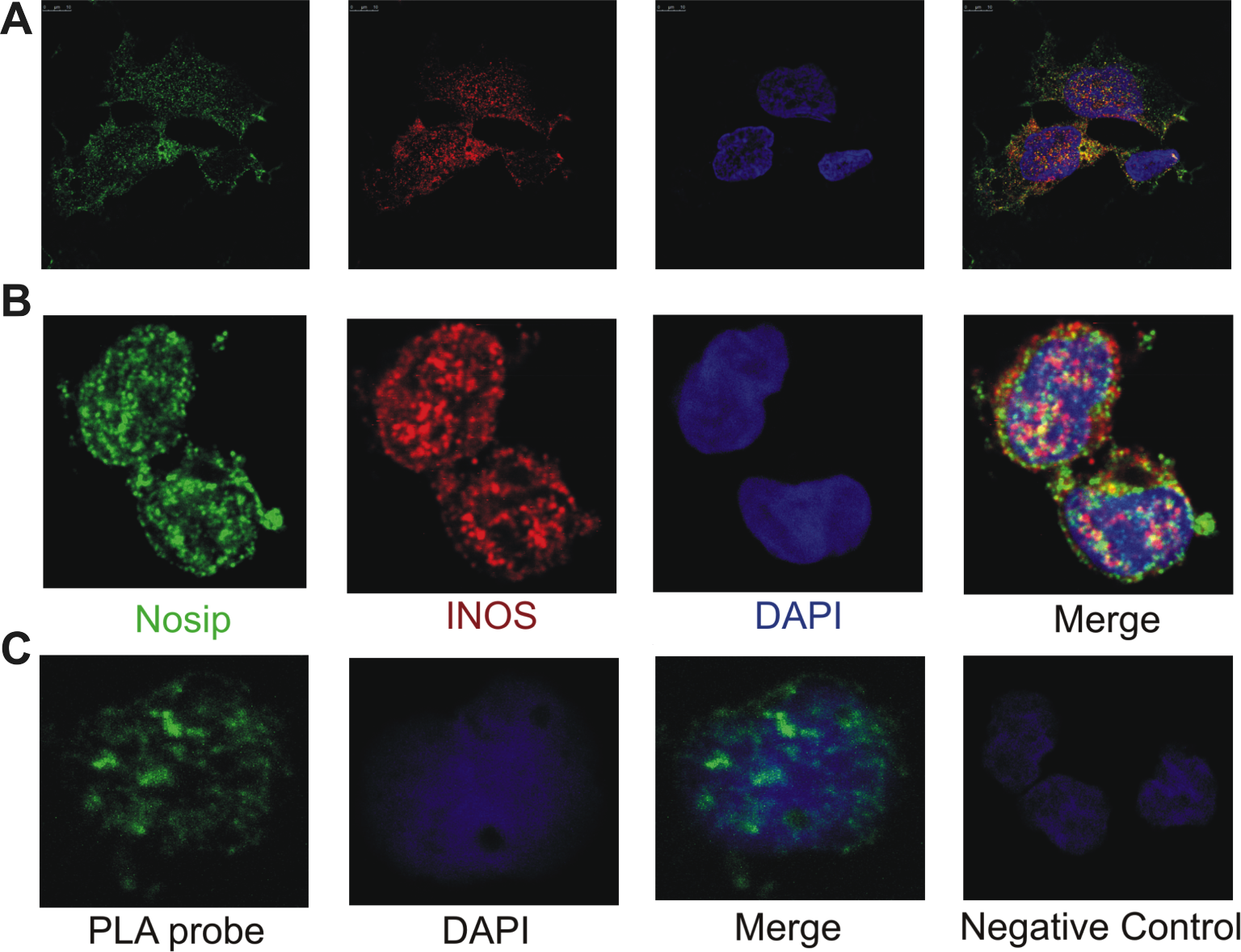
Cellular localization of iNOS and NOSIP. Cellular localization of iNOS and NOSIP in monocytic cell – line THP – 1 **(A)** and primary human monocytes **(B)** showing co - localization of the two proteins. **(C)** In situ proximity ligation assay demonstrating diresct physical interaction between iNOS and NOSIP in primary human monocytes. Each speckle corresponds to one interacting pair. Negative control cells with only one primary antibody shows no background staining.

**Figure S2.**
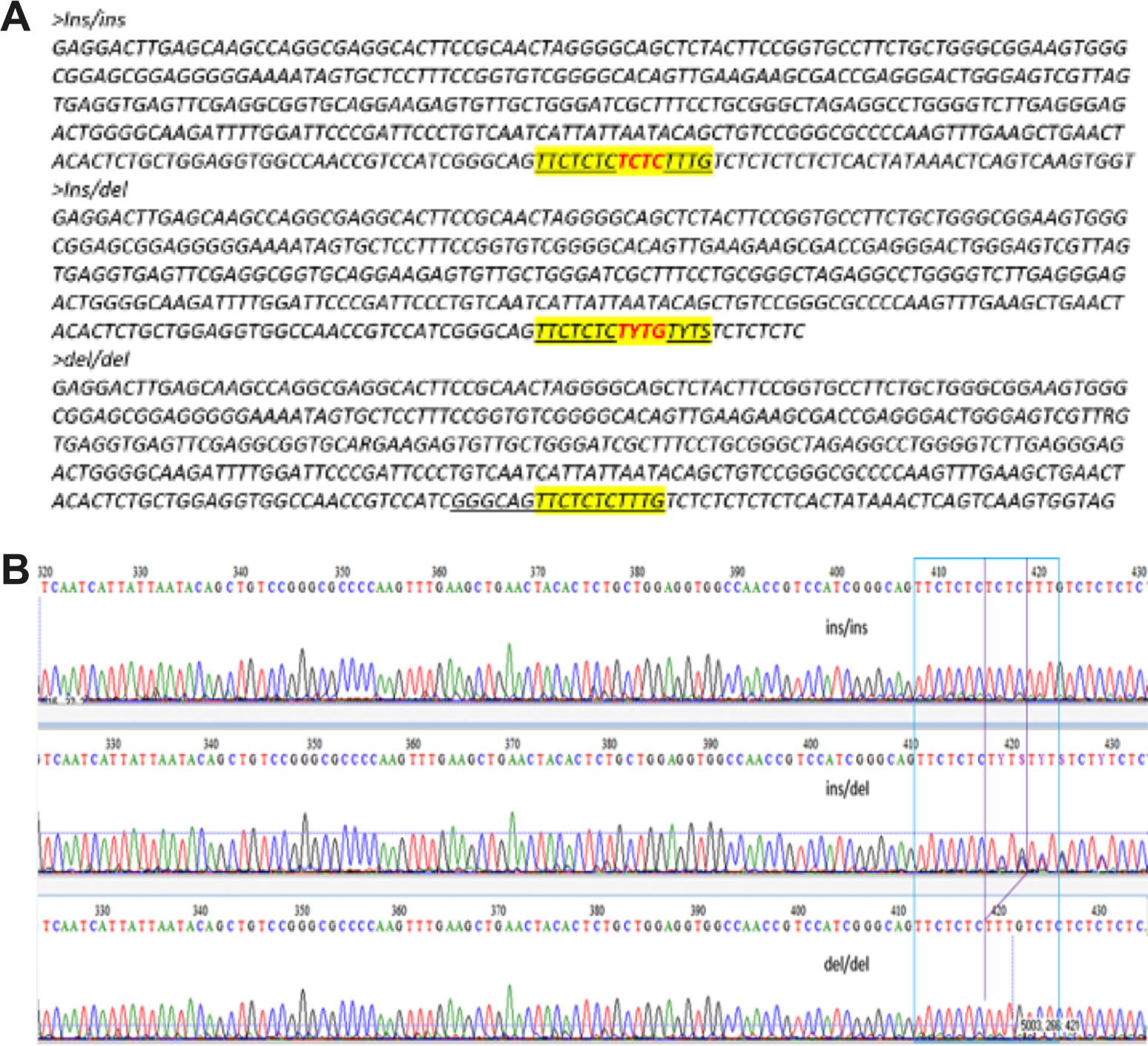
Sequence analysis of NOSIP polymorphism. (A) Representative sample sequencing showing the indel region in the sequence (ins/ins, ins/del and del/del region). (B) Representative chromatogram showing the NOSIP indel regions.

**Figure S3.**
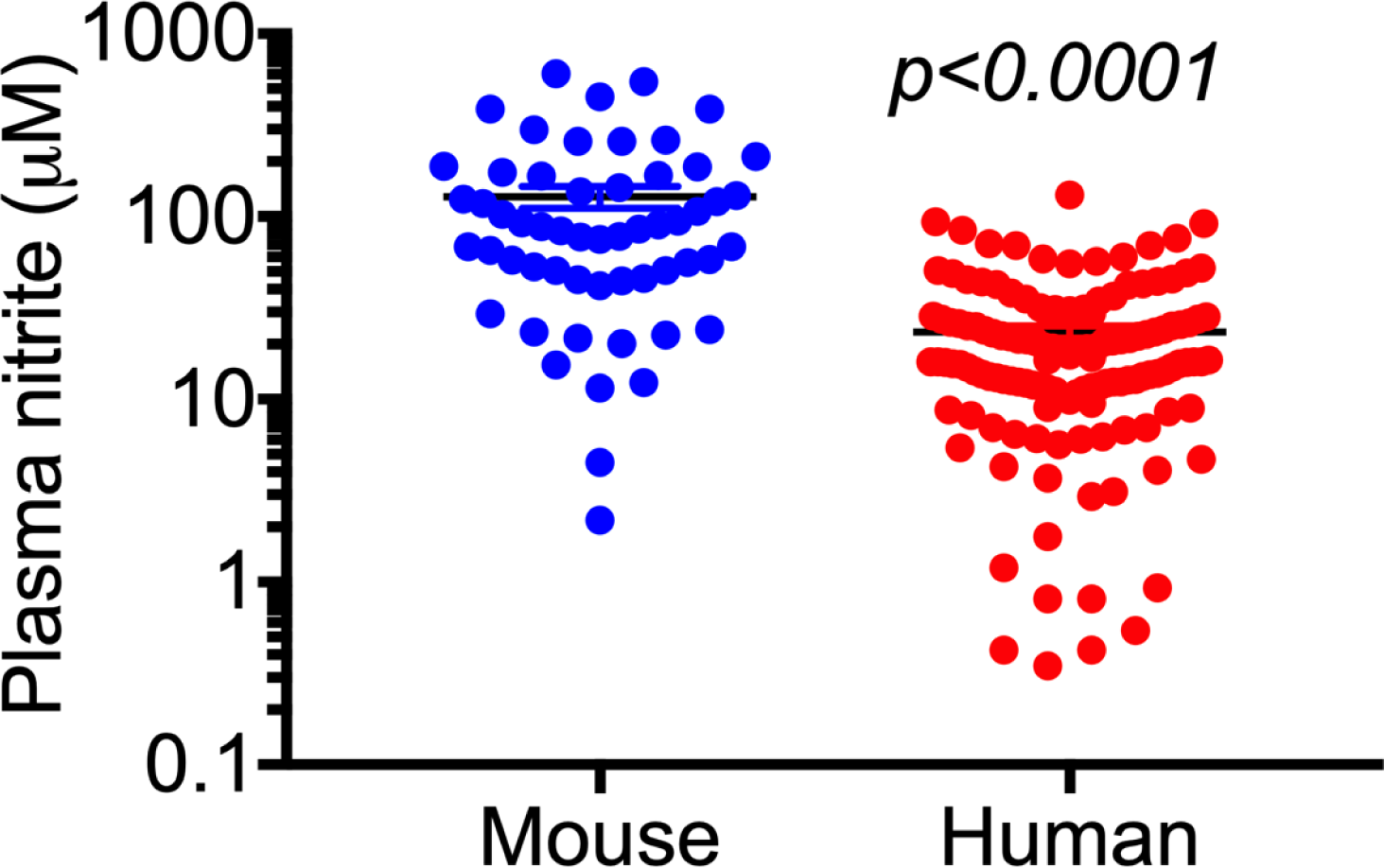
Comparison of plasma nitrite between mouse endotoxemia and human sepsis. Difference in nitrite levels between plasms of human sepsis patients (n = 137) and mice injected with LPS (n = 56). The graph shows overall plasma nitrite levels in the human study cohort that consisted of different categories based on clinical severity and the mice cohort that had two different doses LPS treatment (5 mg/kg and 35 mg/kg) at four different time points (2, 4, 8, and 16 hours). Statistical significance was measured by an unpaired Student’s t-test with Welch’s correction.

**Figure S4.**
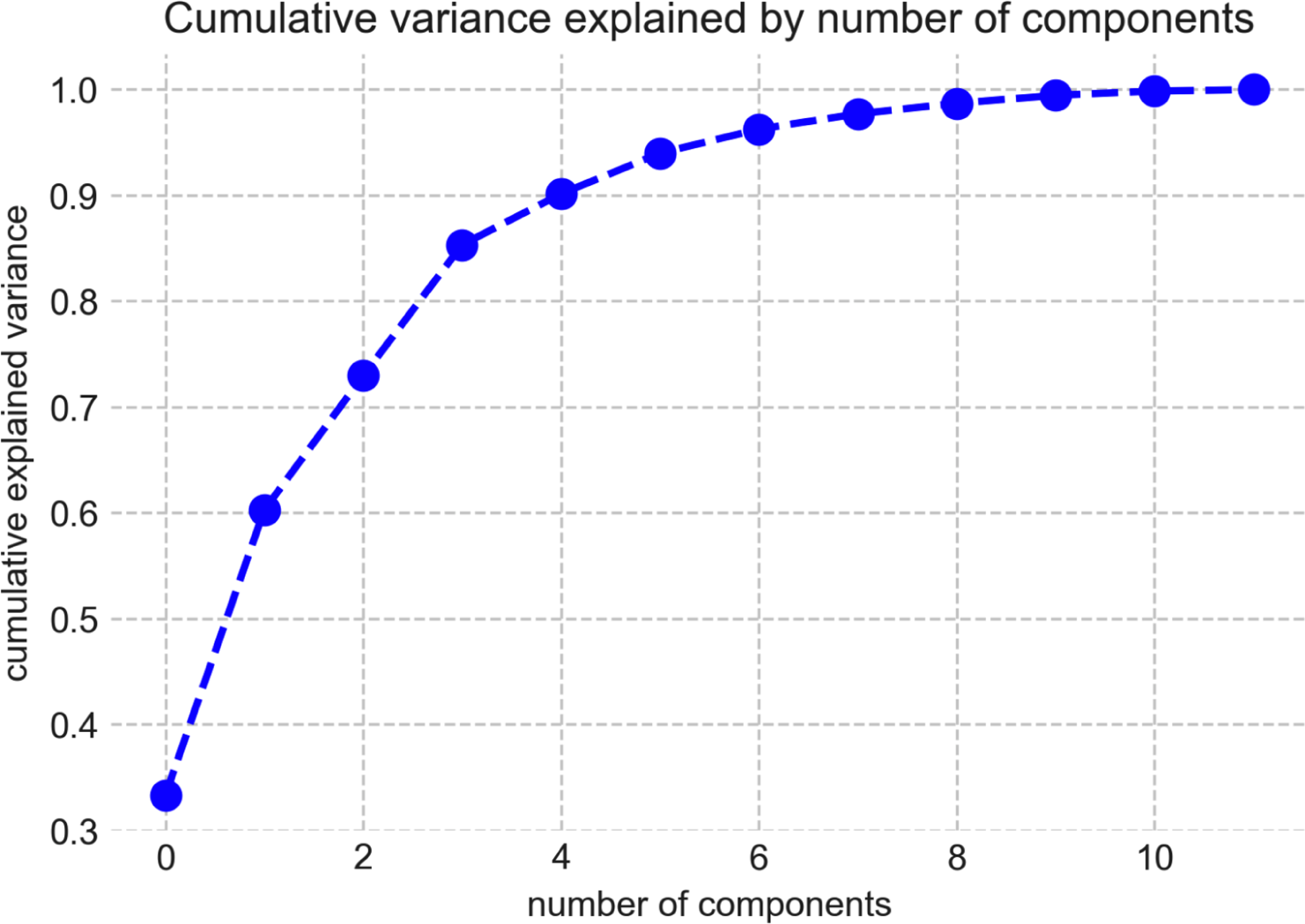
Elbow plot of principal components. Elbow plot showing the contribution of each principal component to the overall variance observed in the data.

